# Valproic acid stimulates myogenesis in pluripotent stem cell–derived mesodermal progenitors in a Notch-dependent manner

**DOI:** 10.1101/2020.12.16.423053

**Authors:** Natacha Breuls, Nefele Giarratana, Laura Yedigaryan, Paolo Carai, Stephane Heymans, Adrian Ranga, Christophe M. Deroose, Maurilio Sampaolesi

**Affiliations:** Laboratory of Translational Cardiomyology, Department of Development and Regeneration, Stem Cell Research Institute, KU Leuven, 3000 Leuven, Belgium; CARIM School for Cardiovascular Diseases, Department of Cardiology, Maastricht University, 6229 ER Maastricht, the Netherlands; Department of Cardiovascular Sciences, KU Leuven, 3000 Leuven, Belgium; Laboratory of Bioengineering and Morphogenesis, Biomechanics Section, Department of Mechanical Engineering, KU Leuven, Leuven, Belgium; Department of Nuclear Medicine, University Hospital KU Leuven, Leuven, Belgium; Human Anatomy Unit, Department of Public Health, Experimental and Forensic Medicine, University of Pavia, 27100 Pavia, Italy

## Abstract

Muscular dystrophies are debilitating neuromuscular disorders for which no cure exists. As this disorder affects both cardiac and skeletal muscle, patients would benefit from a cellular therapy that can simultaneously regenerate both tissues. The current protocol to derive bipotent mesodermal progenitors which can differentiate into cardiac and skeletal muscle relies on the spontaneous formation of embryoid bodies, thereby hampering further clinical translation. Additionally, as skeletal muscle is the largest organ in the human body, a high myogenic potential is necessary for successful regeneration. Here, we have optimized a protocol to generate chemically defined induced pluripotent stem cell-derived mesodermal progenitors (cdMiPs). We demonstrate that these cells contribute to myotube formation and differentiate into cardiomyocytes, both *in vitro* and *in vivo.* Furthermore, the addition of valproic acid, a clinically approved small molecule, increases the potential of the cdMiPs to contribute to myotube formation without compromising their ability to differentiate towards cardiomyocytes. This effect is mediated through the activation of the Notch signaling pathway. Taken together, these results constitute a novel approach to generate mesodermal progenitors with enhanced myogenic potential using clinically approved reagents, which opens the door to new therapeutic solutions in the treatment of muscular dystrophy.

## Introduction

Muscular dystrophies (MDs) are a debilitating group of muscle disorders for which no cure exists. Patients not only suffer from progressive deterioration of the skeletal muscles, leading to a decreased walking ability, but also develop severe cardiomyopathy later on *(1)*. Therefore, these patients would benefit from a cellular therapy capable of targeting both striated muscle types. Induced pluripotent stem cells (iPSCs) have a high proliferative capacity and can differentiate towards all three embryonic lineages. These characteristics make iPSCs an attractive cell type for regenerative medicine. Furthermore, through differentiation towards mesoderm, bipotent stem cells can be generated. Recently, a first protocol was developed to generate human iPSC-derived mesodermal progenitors (MiPs) by means of embryoid body (EB) formation and subsequent sorting for CD44, CD140A and CD140B *(2, 3)*. The obtained MiPs had the *in vitro* and *in vivo* ability to differentiate towards both the cardiac and skeletal muscle lineages. Nevertheless, improvements are needed for further translation. Firstly, the use of EBs is very laborious and heterogeneous, making it necessary to perform subsequent sorting, thus limiting their translatability. Furthermore, MiPs derived from fibroblasts had a much lower myogenic potential than those derived from myogenic progenitors such as mesoangioblasts (MABs) *(2)*. As fibroblasts are easier to obtain and thus a more attractive source to generate iPSCs, strategies need to be developed to overcome this difference.

The Notch signaling pathway is a highly conserved communication system known to be important for myogenesis during embryonic development and postnatal regeneration *(4, 5)*. Furthermore, stimulation of Delta-like 1 (Dll1)/Notch1 has shown to stimulate the regenerative capacity of MABs *(6)*. In previous studies, the Notch1 pathway could be activated through the addition of valproic acid (VPA) *(7)*. VPA is a small molecule known for its function as a histone deacetylase inhibitor (HDACi). Interestingly, VPA has shown to stimulate myogenesis in both myogenic progenitors and iPSCs *(8, 9)*.

In this study, we aimed to derive human MiPs through a fully chemically defined monolayer approach, thereby enhancing their clinical translatability. Furthermore, we asked whether the Notch signaling pathway can be used to stimulate the myogenic potential of these newly developed chemically defined MiPs (cdMiPs). In this context, we will investigate if VPA can be used to stimulate Notch1 and thus lead to an increased myogenic potential in a translatable manner.

## Results

### Chemically defined mesodermal progenitors have the ability to differentiate into the myogenic and cardiac lineages

In order to obtain cdMiPs, human iPSCs were differentiated for four days following a serum-free monolayer approach (Fig. 1A). During the differentiation, the cells gradually lost pluripotency markers *OCT4* and *NANOG,* while early mesodermal markers such as *BRACH, MSGN1* and *TBX6* were upregulated after two days of CHIR99021 exposure (Fig. 1B). From day 2 onwards, markers for both paraxial as well as lateral plate mesoderm were upregulated (Fig. 1C). Apart from *CXCR4*, ecto- and endodermal markers stayed more stable over time (Figure 1D).

**Fig. 1.**
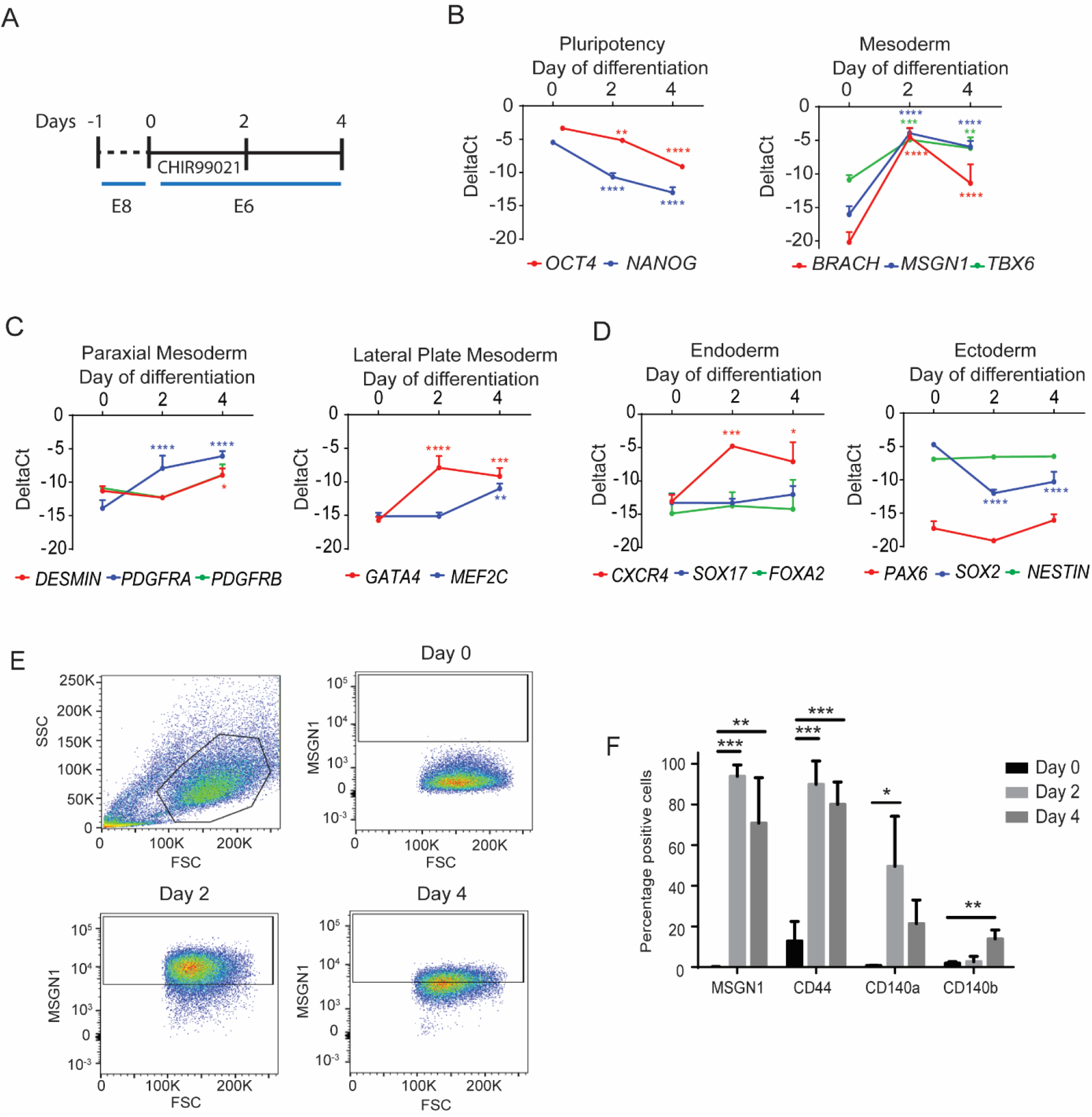
Mesodermal progenitors can be derived trough a chemically defined monolayer approach. (**A**) Schematic overview of the protocol to differentiate induced pluripotent stem cells towards mesodermal progenitors. (**B-D**) Gene expression of markers for pluripotency, early and late mesoderm, ectoderm and endoderm during mesoderm differentiation. (**E**) FACS analysis of Mesogenin1 (MSGN1) during differentiation using a MSGN1-2A-Venus reporter line. (**F**) Percentage of MSGN1, CD44 and CD140A/B positive cells during differentiation, based on FACS analysis. *P<0.05; **P<0.01; ***P<0.001; ****P<0.0001. n=3-4.

Using a human MSGN1-2A-*Venus* iPSC line, 93.9±4.6% of the cells were positive MSGN1^+^ after two days of differentiation, correlated to a strong mesoderm induction (Fig. 1E) *(10)*. Furthermore, looking at the markers previously used to isolate MiPs, a higher percentage of CD44, CD140A and CD140B-positive cells could be observed from day 2 onwards, increased at this stage of differentiation (Fig. 1F).

To confirm the mesodermal nature of our cdMiPs, the cells were differentiated towards skeletal muscle, smooth muscle and osteogenic lineages. After differentiation, the cdMiPs were positive for myosin heavy chain (MyHC), alpha smooth muscle actin (SMA) and alizarin red S, respectively (Fig. 2A-C). To check whether the cells have the potential to contribute to the myogenic and cardiac lineages, cdMiPs were put in co-culture with either C2C12 myoblasts or rat cardiomyocytes. After subsequent differentiation, the lamin A/C-positive cdMiPs expressed either MyHC (6.97±6.19%) or cardiac MyHC (11.72±14.31%) after co-culture with C2C12 myoblasts or rat cardiomyocytes, respectively (Fig. 2D and E). In general, we were able to produce cdMiPs that have the ability to differentiate into multiple mesodermal lineages.

**Fig. 2.**
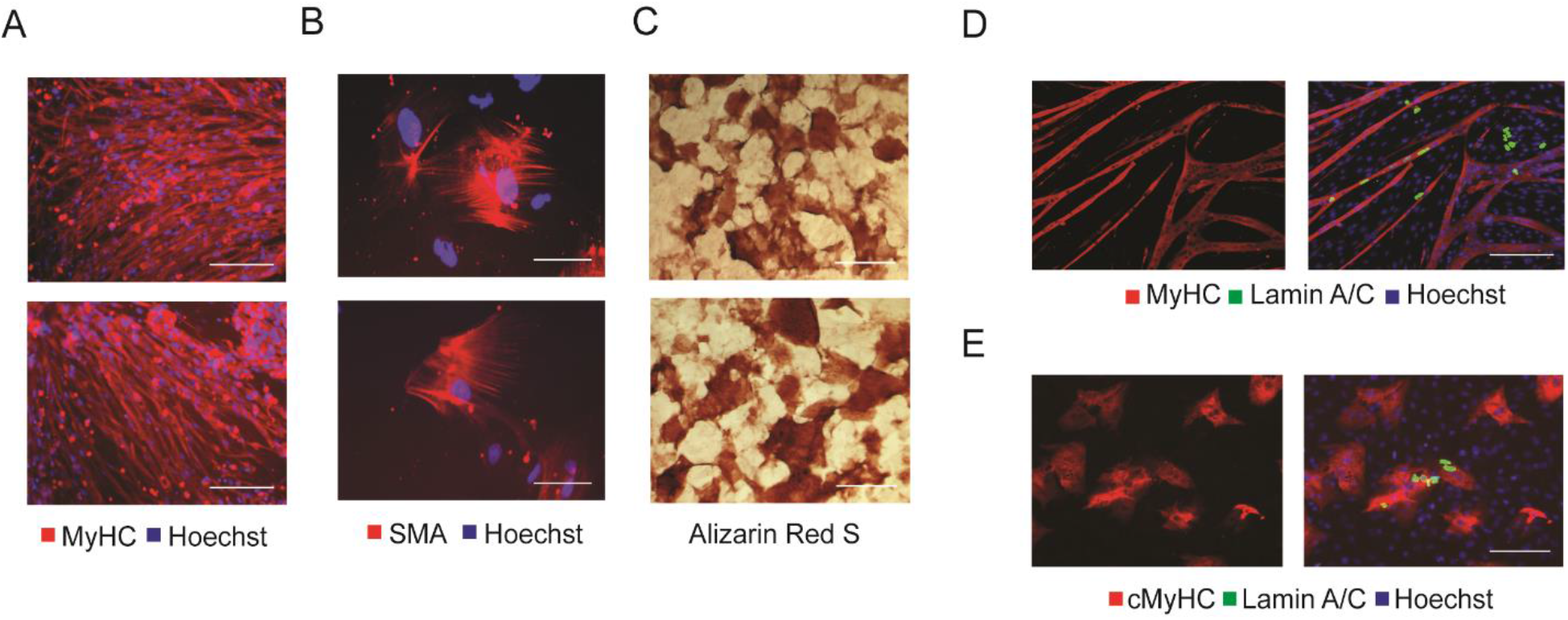
Mesodermal progenitors have the potential to differentiate towards multiple mesodermal lineages and contribute to in vitro cardiac and skeletal muscle formation. (**A**) Immunofluorescence of MyHC (red) after myogenic differentiation. (**B**) Immunofluorescence after smooth muscle differentiation (SMA). (**C**) Alizarin red S staining after osteogenic differentiation of the mesodermal progenitors. (**D-E**) Immunofluorescence analysis of lamin A/C-positive mesodermal progenitors (green) in co-culture with either C2C12 myoblasts (**D**) or rat cardiomyocytes (**E**). Scale bars: 100 μm.

### Mesodermal progenitors show long-term survival and engraftment upon serum exposure

In order to address the *in vivo* functionality of the cdMiPs, a multifunctional imaging reporter gene construct was integrated via zinc finger nuclease (ZFN)-targeting, downstream of exon 1 of the gene phosphatase 1 regulatory subunit 12C *(PPP1R12C)* gene, known as the safe-harbor adeno-associated virus site 1 *(AAVS1)* locus *(11)*. The reporter genes were driven by the hybrid CAGGS promoter consisting of an early cytomegalo virus enhancer fused to a chicken beta-actin promoter. The reporter genes used were enhanced green fluorescent protein (eGFP) for immunofluorescence, firefly luciferase (Fluc) for bioluminescence imaging (BLI) and human sodium-iodide symporter (hNIS) for positron emission tomography (PET) (Fig. S1A). Here, the hNIS reporter gene has the benefit of being used already in a clinical setting for tumor detection, and can thus be used in patients for stem cell tracking *(12)*. A specific integration of the construct into the AAVS1 locus was confirmed via a junction assay and Southern blot analysis (Fig. S1B and C). After the integration, the cells maintained their expression of pluripotency factors *OCT4* and *NANOG* and showed expression of the hNIS reporter gene, Solute Carrier Family 5 Member 5 *(SLC5A5}* (Fig. S1D). Functional Fluc was validated via BLI (Fig S1E) and eGFP could be detected via immunofluorescence (Figure S1F). Furthermore, the uptake of pertechnetate (^99m^TcO_4_^-^) confirmed functionality of the hNIS. This uptake was completely abolished upon administration of sodium perchlorate (NaClO_4_), a specific hNIS blocker, confirming specific uptake through the hNIS (Fig. S1G).

Once the reporter construct was integrated, 1.5 x 10^6^ cdMiPs were injected into the hind limbs of beta-sarcoglycan-null *(Sgcb^-/-^)/* common gamma knockout *(Rag2γc^-/-^)* mice. After injection, the BLI signal was lost after two days, indicating a low survival rate (Fig. 3A). In order to prepare the cells better for the *in vivo* environment, fetal bovine serum (FBS) was added to the cdMiPs from day 3-4 of differentiation. Pre-exposure of the cells to 10% FBS led to a higher signal right after injection compared to the untreated cdMiPs. However, this signal was lost seven days postinjection (Fig. 3B). When the concentration of FBS was increased to 20%, the cells showed longterm survivability (Fig. 3C). After an initial drop, the BLI signal could be maintained for up to 28 days post-injection (Fig. 3D). Although exposure to serum can introduce heterogeneity in the culture, the addition of 20% FBS did not lead to any changes in the gene expression of key meso, ecto- and endodermal markers (Fig. S2).

**Fig. 3.**
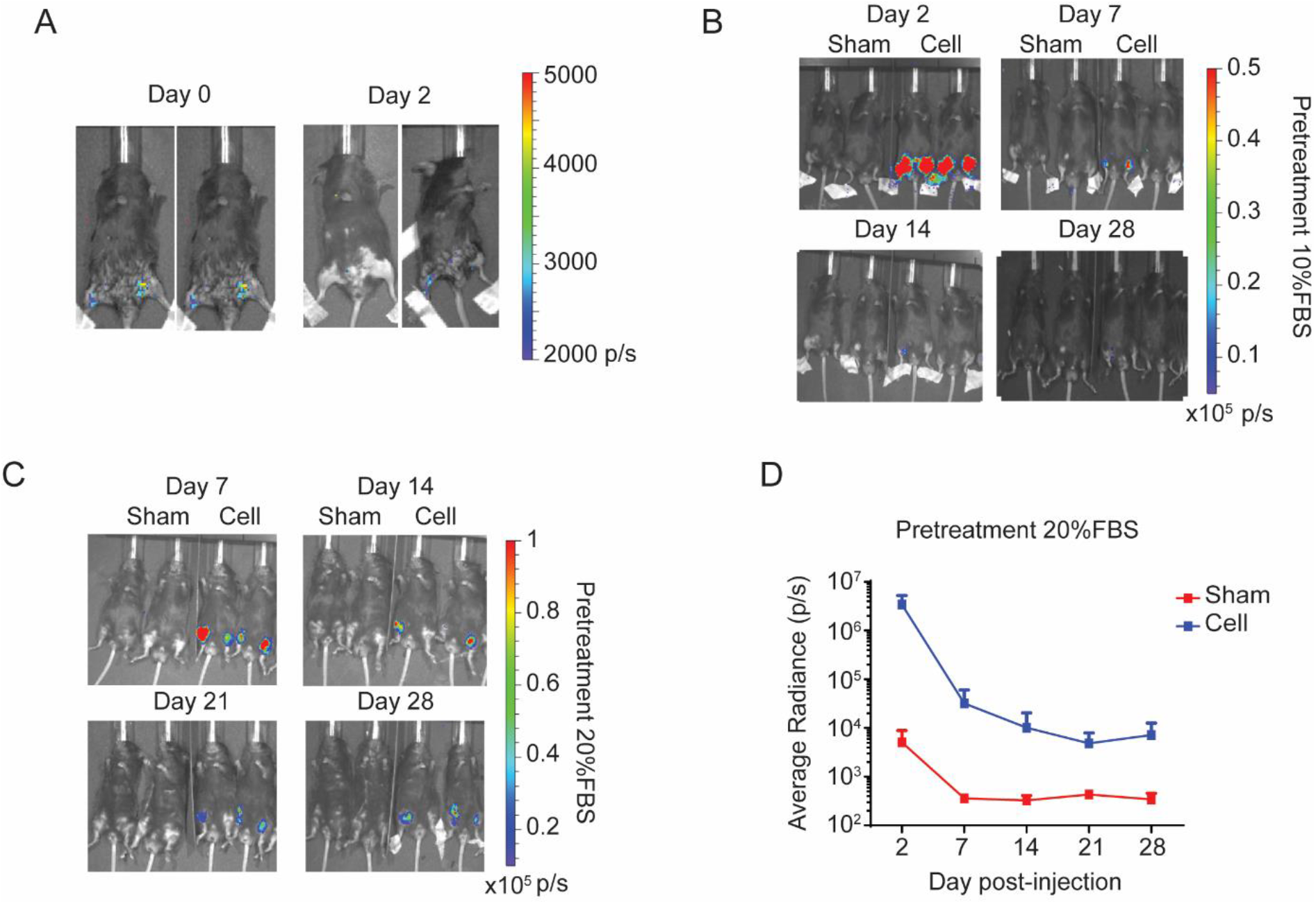
Serum pre-treatment of mesodermal progenitors leads to long-term survivability and stable engraftment in the skeletal muscle. (**A**) Representative bioluminescent images of beta-sarcoglycan-null *(Sgcb^-/-^)/* common gamma knockout *(Rag2γc^-/-^)* mice injected in both hind limbs with 1.5 x 10^6^ mesodermal progenitors (cdMiPs). (**B**) Representative bioluminescent images of *Sgcb^-/-^/Rag2γc^-/-^* mice injected with saline (sham) or 1.5 x 10^6^ cdMiPs treated with 10% serum (cell) in the hind limbs. Images were taken 2-, 7-, 14- and 28-days post-injection. (**C**) Representative bioluminescent images of Sgcb^-/-^/Rag2γc^-/-^ mice injected with saline (sham) or 1.5 x 10^6^ cdMiPs pre-exposed to 20% serum (cell) in the hind limbs. Images were taken 7-, 14-, 21- and 28-days post-injection. (**D**) Quantification of the BLI signal seen in (**C**) (n=3).

After 28 days, GFP^+^ cdMiPs could be detected within the muscle fibers of the skeletal muscle. Besides their fusion into existing myofibers, individual GFP^+^ LAMININ^+^ cells were detected, indicating differentiation of the cdMiPs into the myogenic lineage (Fig. 4A). To investigate whether the cdMiPs have the ability to engraft in the heart, 1.5 x 10^6^ cdMiPs were injected in the wall of the left ventricle of *Sgcb^-/-^/Rag2γc^-/-^* mice. After injection, individual GFP^+^ sarcomeric alpha-actinin^+^ cells were detected, indicating differentiation of the cdMiPs towards the cardiac lineage (Fig. 4B). These data reveal that the cdMiPs can engraft in both cardiac and skeletal muscle tissues. However, prior exposure to serum is necessary to guarantee long-term survival upon injection.

**Fig. 4.**
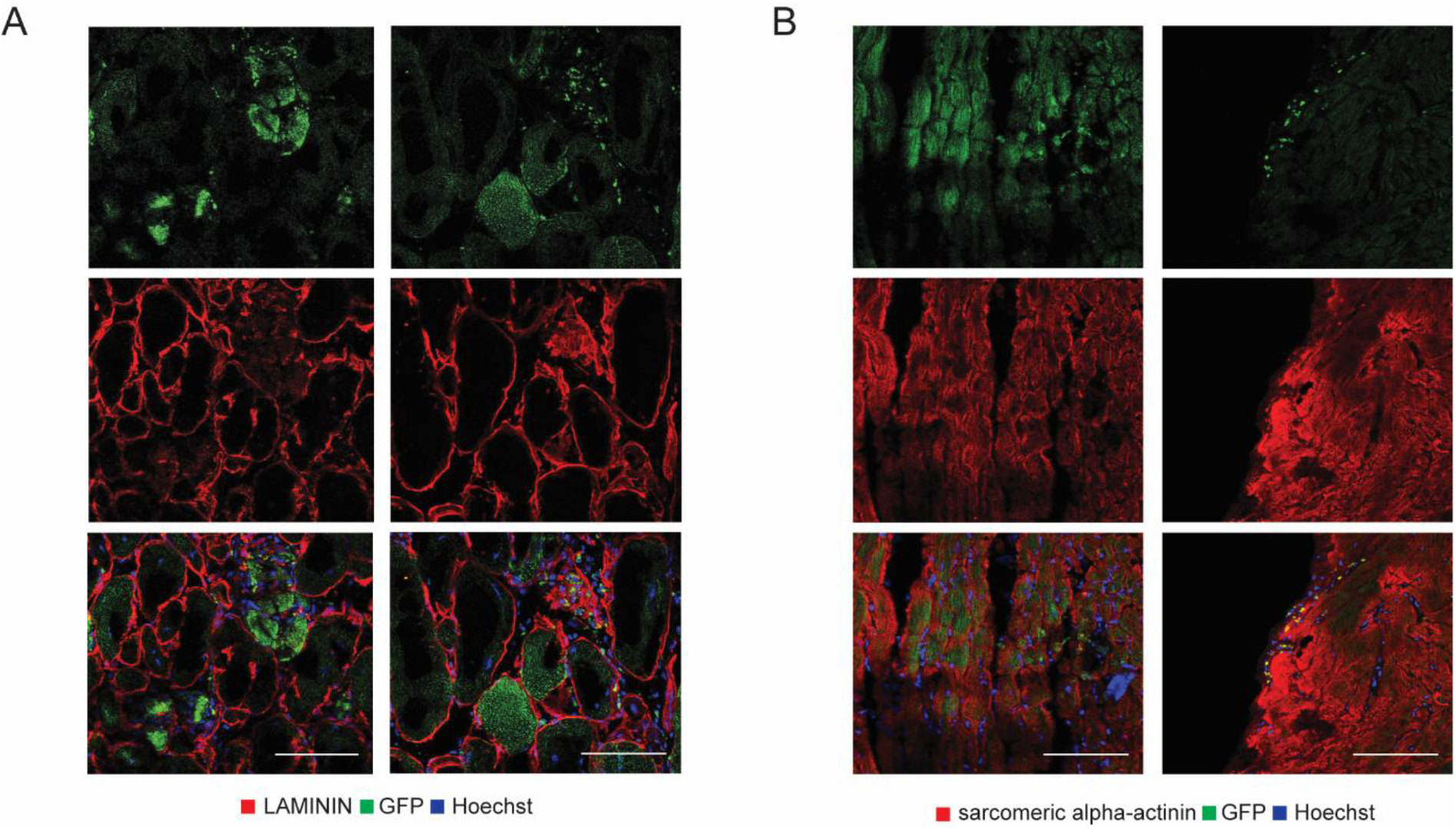
Mesodermal progenitors have the ability to engraft into the cardiac tissue of dystrophic mice. (**A**) Immunofluorescent images of the GFP^+^ cdMiPs present in the skeletal muscle 28 days after injection. (**B**) Immunofluorescence of the GFP^+^ cdMiPs present in the heart, 28 days after injection. Scale bars: 100 μm.

### Notch activation through Dll1 increases the myogenic potential of cdMiPs

Recently, our group identified that induction of the Dll1/Notch1 signaling pathway could enhance the myogenic regenerative capacity of MABs *(6, 13)*. MABs resemble the generated cdMiPs, as they both can differentiate towards multiple mesodermal lineages and show the expression of CD44, CD140B and CD140B (Fig. 1). Therefore, we wanted to determine if induction of the Notch pathway through Dll1 could improve the myogenic potential of cdMiPs.

In order to stimulate the Notch signaling pathway, a construct containing a doxycycline (dox)-inducible *DLL1* was inserted into a human embryonic stem cell (ESC) master cell line by means of recombinase-mediated cassette exchange (RMCE) (Fig. 5A) *(14)*. After integration, a higher amount of *DLL1* became apparent after four days of dox treatment (Fig. 5B and C). Adding dox during the entire mesodermal differentiation led to an increased contribution of these cells to myotubes upon co-culture (Fig. 5D and E). These results suggest that activation of the Notch signaling through Dll1 drives cdMiPs towards the myogenic lineage.

**Fig. 5.**
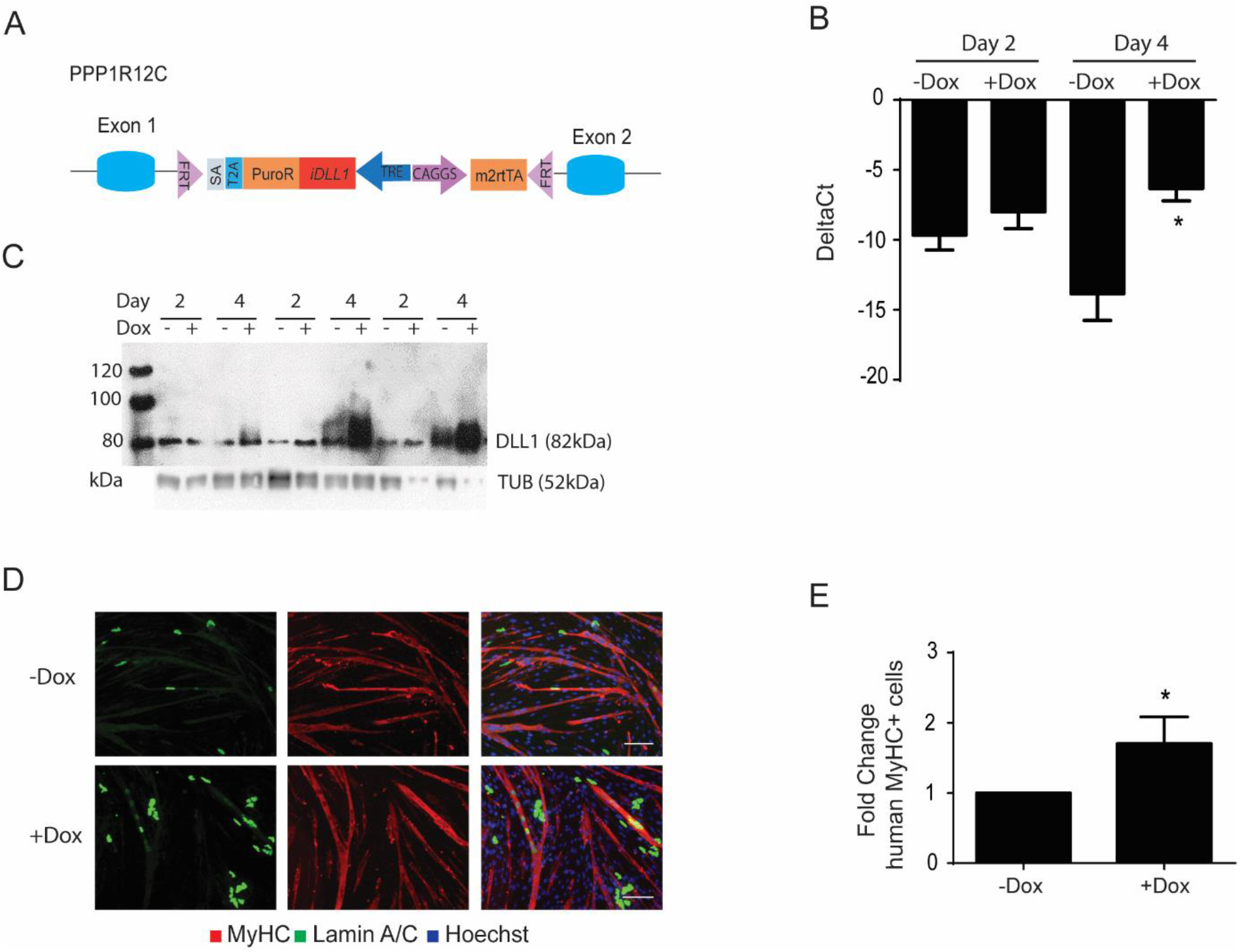
*DLL1* overexpression improves the myogenic propensity of mesodermal progenitors. (**A**) Schematic overview of the doxycycline (dox)-inducible DLL1 construct flipped into an embryonic stem cell (ESC) master line. (**B**) Quantitative PCR for *DLL1* after exposure of the generated ESC line to dox for two or four days. (**C**) Western blot for DLL1 after exposure of the generated ESC line to dox for two or four days. Tubulin (TUB) was used as a housekeeping protein. (**D**) Immunofluorescence analysis of lamin A/C-positive mesodermal progenitors, with or without dox, (green) in co-culture with C2C12 myoblasts (n=3). Scale bars: 100 μm. *P<0.05.

### Valproic acid relies on the Notch signaling pathway to stimulate myogenesis

To use the beneficial effect of Notch activation, a small molecule is needed to induce Notch signaling in a manner that allows future clinical translation. Previously studies already showed that VPA can stimulate Notch1 signaling pathway *(7)*. Furthermore, VPA has shown to enhance the myogenic capacity of adult stem cells and to induce myogenesis directly in iPSCs *(8, 9)*. To test its effect on the cdMiPs, 1mM VPA was added from day 2-4 of differentiation. Afterwards, the cells were put in co-culture with either C2C12 myoblasts or rat cardiomyocytes (Fig. 6). Upon VPA treatment, ~3-fold more lamin A/C-positive nuclei were detected in MyHC^+^ myotubes, suggesting a beneficial effect of VPA on the myogenic capacity of the cdMiPs (Fig. 6A and B). Additionally, no decrease was detected in the amount of lamin A/C-positive cardiomyocytes after exposure to VPA (Fig. 6C and D). These results suggest that VPA stimulates the myogenic capacity of cdMiPs without interfering with their ability to form cardiomyocytes.

**Fig. 6.**
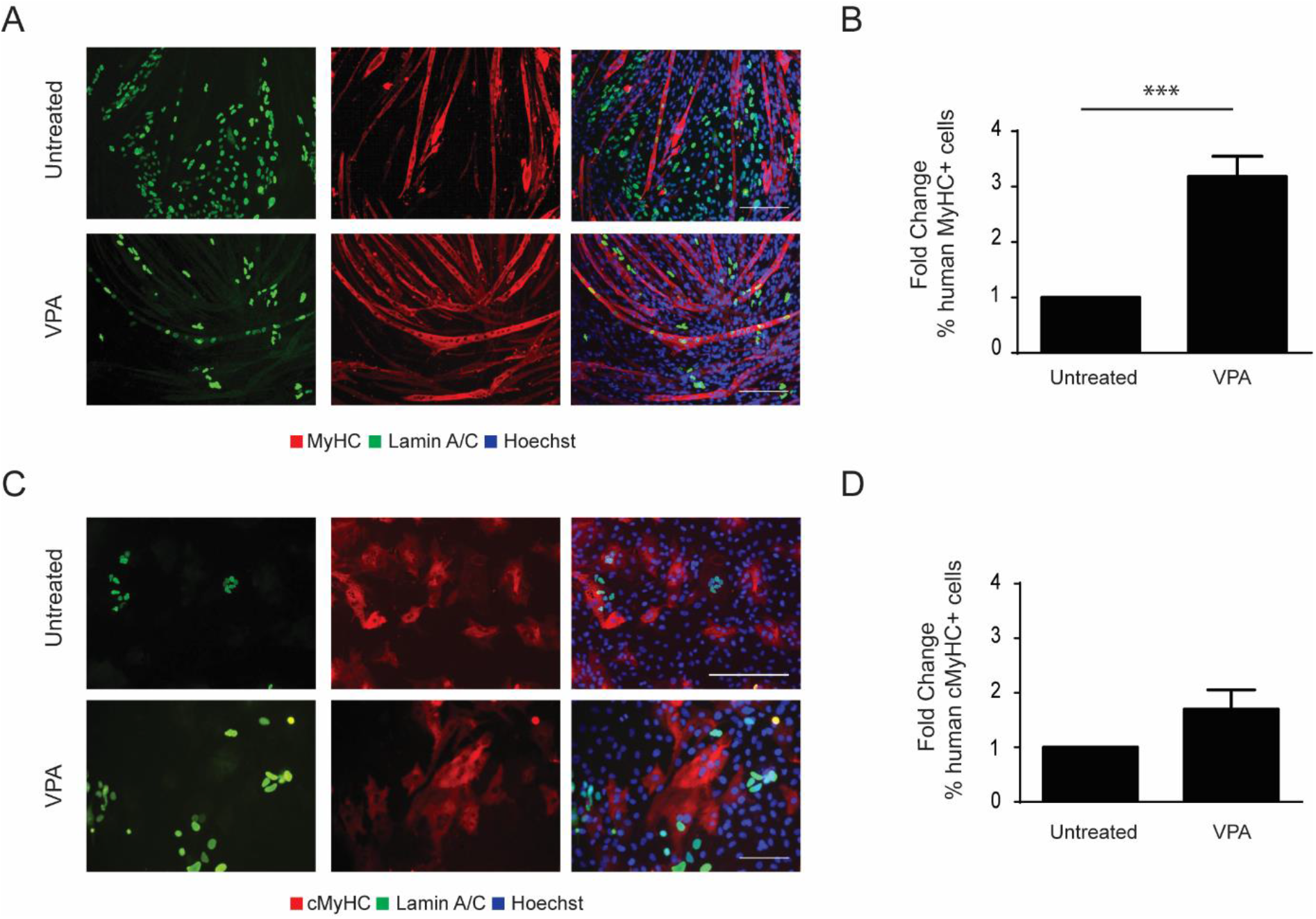
Valproic acid increases the myogenic potential of mesodermal progenitors while maintaining the cardiogenic potential. (**A**) Immunofluorescence of lamin A/C-positive mesodermal progenitors (green), with or without exposure to valproic acid (VPA) in co-culture with C2C12 myoblasts. (**B**) Amount of lamin A/C-positive mesodermal progenitors positive for myosin heavy chain (MyHC), compared to the untreated condition (n=4). (**C**) Immunofluorescence of lamin A/C-positive mesodermal progenitors (green), with or without exposure to VPA, in co-culture with rat cardiomyocytes. (**D**) Amount of lamin A/C-positive cells positive for cardiac myosin heavy chain (cMyHC), compared to the untreated condition (n=3). Scale bar: 100μm. ***P<0.001.

To check whether VPA has the ability to stimulate the Notch1 signaling pathway in cdMiPs, the presence of the Notch intracellular domain (NICD) was checked after 0, 6, 12, 24 and 48 hours of VPA exposure. After 24 hours, an increased amount of the NICD was detected, indicating activation of the Notch signaling pathway. However, after 48 hours of VPA exposure, a Notch1 inhibition could be observed (Fig. 7A and B).

**Fig. 7.**
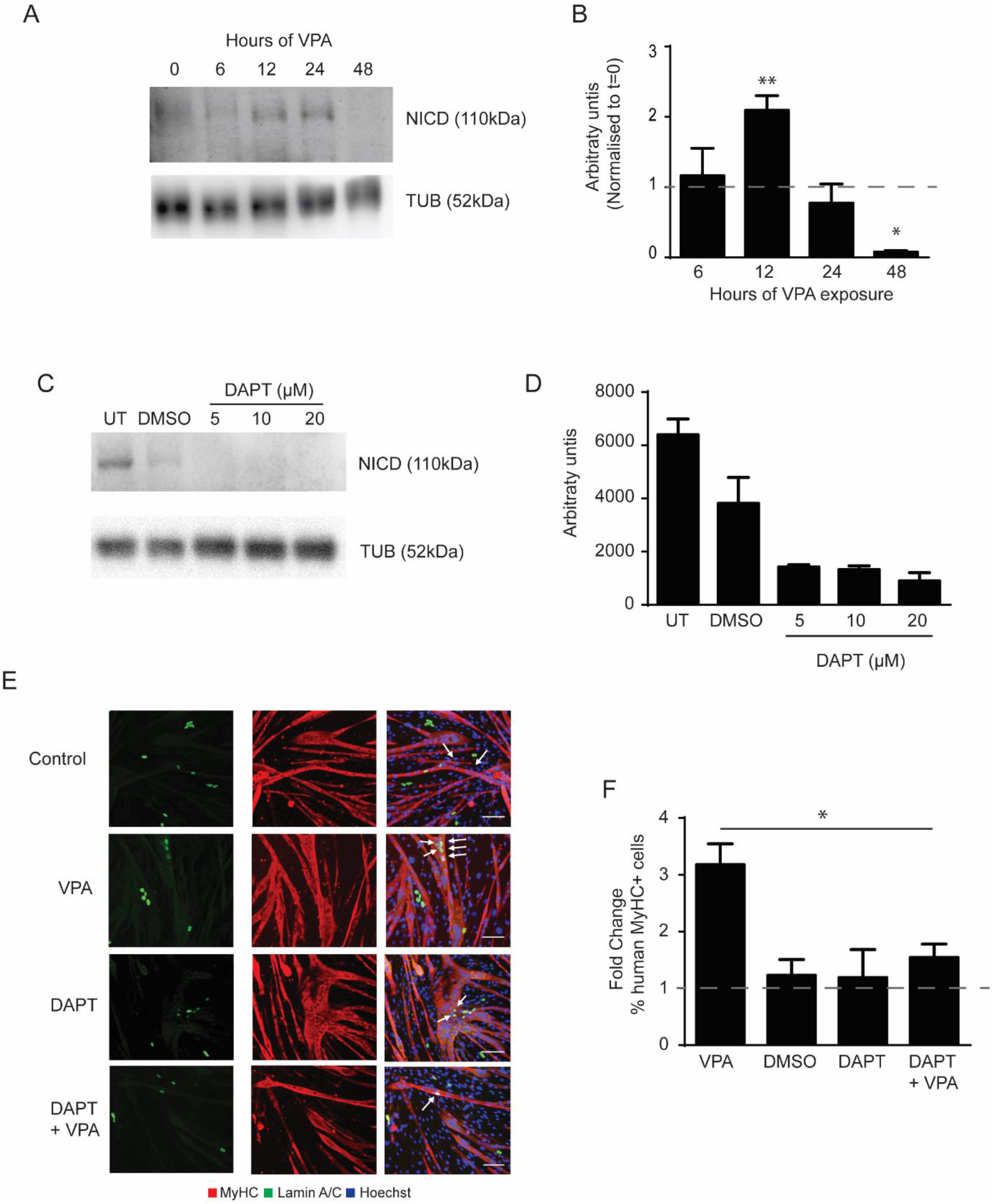
Activation of the Notch signaling is required for valproic acid to stimulate myogenesis. (**A**) Representative western blot for the Notch1 intracellular domain (NICD) of mesodermal progenitors exposed to valproic acid (VPA) for 0, 6, 12, 24 and 48 hours. Tubulin (TUB) was used for normalization. (**B**) Quantification of (**A**) normalized to no exposure to VPA. *P<0.05; **P<0.01. n=3. (**C**) Representative western blot for the NICD of mesodermal progenitors exposed to dimethyl sulfoxide (DMSO) or 5/10/20 μM of DAPT from day 0-4 of differentiation. TUB was used for normalization. (**D**) Quantification of (**C**). n=2. (**E**) Immunofluorescence of lamin A/C-positive mesodermal progenitors (green), untreated or with exposure to VPA, DMSO (control), DAPT or DAPT+VPA, in co-culture with C2C12 myoblasts. (**F**) Fold change of lamin A/C-positive cells, positive for myosin heavy chain (MyHC) compared to the appropriate control (untreated or DMSO). n=3. Scale bar: 100 μm. *P<0.05.

In order to investigate whether an initial increase in Notch is necessary for the pro-myogenic effect of VPA, γ-secretase inhibitor DAPT was added from day 0-4 of differentiation to inhibit the Notch signaling pathway. DAPT prevents cleavage of the Notch receptor and thus prevents the release of the NICD. Addition of 5-20 μM DAPT led to a decrease of the NICD (Fig. 7C and D). When DAPT was added during the entire length of the differentiation, no difference could be observed in the amount of lamin A/C^+^ cells in the MyHC^+^ myotubes compared to the dimethyl sulfoxide (DMSO)-treated control. However, when DAPT was added together with VPA, the increase normally seen in lamin A/C^+^ MyHC^+^ cells, is abolished (Fig. 7E and F). Taken together, it seems that the initial activation of the Notch signaling pathway is important for VPA to elicit its pro-myogenic effect.

### Single-cell RNA sequencing shows that valproic acid pushes cell towards a myogenic state

To get more insight on how VPA affects the cdMiPs, single-cell RNA sequencing (scRNAseq) was performed of both untreated and VPA-treated cells using SMART-Seq2 *(15)*. Afterwards, thorough quality control, we retained 227 high-quality cells (Fig. S3A-B). Upon SC3 clustering, the t-distributed stochastic neighborhood embeddings (t-SNE) plot can be divided into three main clusters. The two largest corresponded to the untreated cdMiPs and those exposed to VPA, while the third cluster mainly contained cells in metaphase (Fig. S3C and D). The use of two clusters led to a division that completely depended on the treatment with an even distribution of the cell cycle state, and was used for further analysis (Fig. 8A). We used gene set enrichment analysis to annotate the clusters further with correlations to the ARCHS4 tissues database *(16)*. Here, we found that the gene expression profile of the VPA-treated group correlated more to myoblasts compared to the untreated group (Fig. 8B-C). Some key myogenic genes found to be differentially expressed were platelet-derived growth factor receptor A (*PDGFRA)*, which decreased upon VPA treatment, and *DESMIN* and *CD 82,* which were upregulated upon VPA supplementation (Fig. 8D).

**Fig. 8.**
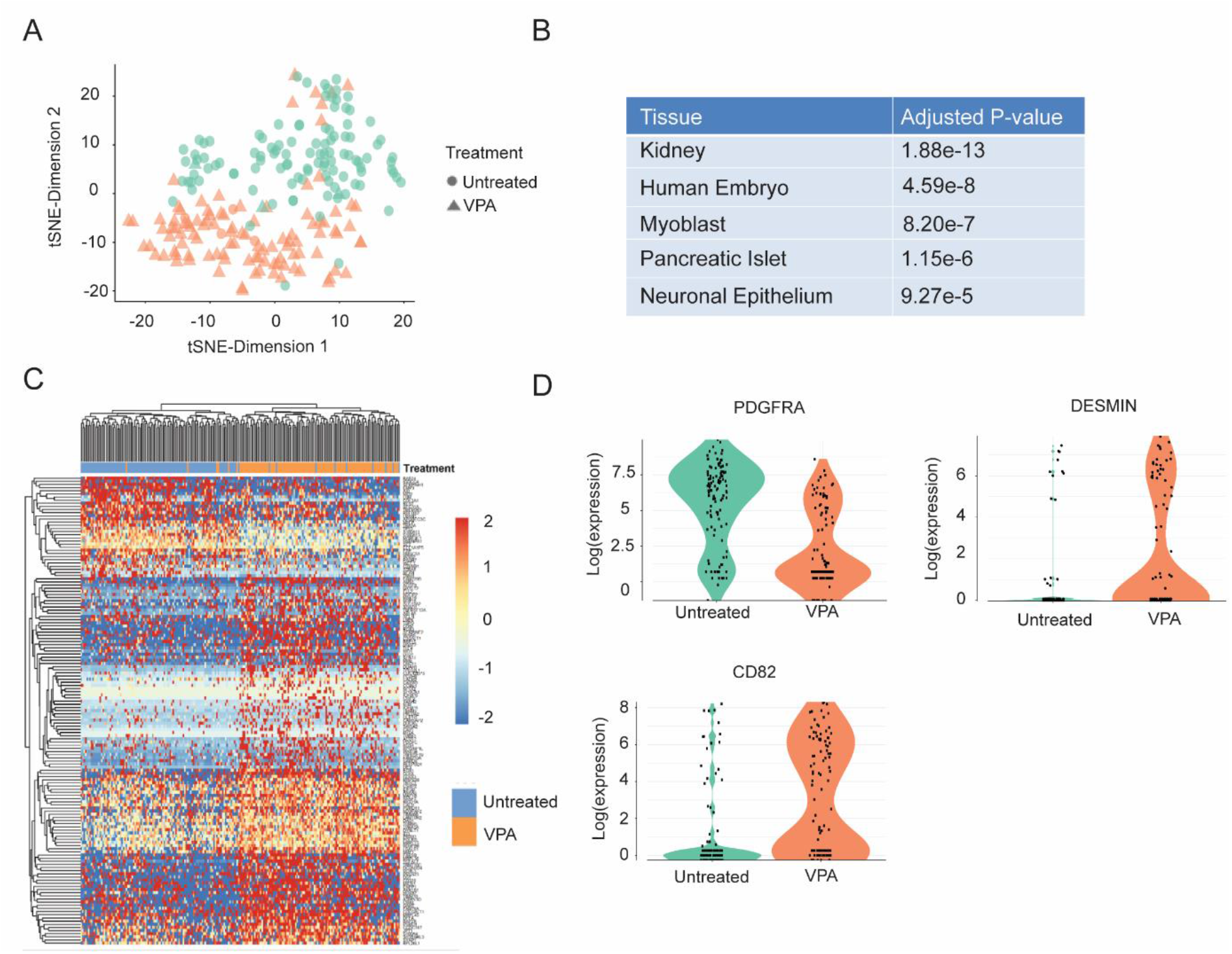
Single-cell RNA sequencing reveals an upregulation of myogenic regulators after valproic acid treatment. (**A**) t-SNE plot colored by k-means clustering (k=2) of 227 cells either untreated (circles) or treated with valproic acid (VPA, triangles). (**B**) Top 5 ARCHS4 significantly enriched in the cluster corresponding to the VPA-treated cells. (**C**) Heatmap of k-means clusters with myogenic-related genes differentially expressed between untreated and VPA-treated mesodermal progenitors. (**D**) Expression of platelet-derived growth factor receptor A (PDGFRA), DESMIN and CD82 in untreated and VPA-treated mesodermal progenitors.

## Discussion

Dual progenitors targeting both skeletal and cardiac muscles are of great interest to target the most debilitating manifestation of MDs. Recently, our laboratory was the first to develop mesodermal progenitors from iPSCs that could successfully differentiate into skeletal and cardiac lineages both *in vitro* and *in vivo (2)*. A recent review provided a blueprint for translational regenerative medicine depicting that successful translation requires manufacturing processes that is reliable and can be scaled while remaining financially viable *(17)*. In this study, we identified a robust approach to generate mesodermal progenitors. By using chemically defined media and differentiating the cells in a monolayer, we provided a more homogeneous method that can be scaled up more easily. Despite the fact that the protocol was performed in a serum-free manner, a 24-hour pre-exposure to FBS was needed to guarantee *in vivo* survival. It has already been shown that serum-deprivation induces apoptosis, suggesting a possible protective effect of serum against cell death *(18, 19)*. The same was found by Kim et al. who showed a low survivability upon transplantation of myogenic progenitors derived from a serum-free protocol *(20)*. The addition of serum for 24 hours did not lead to any major changes in important lineage markers. However, animal products should be avoided, as there is a risk of the presence of non-human pathogens that can induce an unwanted immune response and subsequent cell rejection. Human serum has already been used to replace FBS in the field of mesenchymal stem cells *(21, 22)*. Within the field of MDs, cells were injected into a golden retriever MD model using serum of the recipient dog, leading to significant engraftment *(23, 24)*. This method could be translated into a human setting, where the patients’ own serum can be used to avoid any immune response.

As shown previously for adult stem cells and iPSCs, VPA was able to stimulate the myogenic commitment of our cdMiPs. However, from our scRNAseq data it is clear that the effect of VPA is not limited to the myogenic lineage, as networks from the embryonic, neuronal and pancreatic lineages are also upregulated after VPA-treatment. These findings are also confirmed by literature where VPA has been used to stimulate reprogramming and neuronal differentiation *(25, 26)*. The exact fate that is stimulated thus depends on the environment in which the cells reside. This could pose a problem *in vivo* where the cells might come in contact with a wide range of signals. VPA is known as a very broad HDACi, leading to this wide variety of effects. Pinpointing the exact mechanism through which VPA exerts its effect on myogenesis, might allow us to find a more specific HDACi with less adverse effects. Here, several other HDACi, including the recently approved Givinostat, have shown to stimulate myogenesis in adult stem cells and thus can be considered within this protocol *(27, 28)*. Another strategy could be selecting a subpopulation specifically linked to the myogenic lineage. Here, CD82 could be a candidate as it has been identified recently as a novel marker for human satellite cells *(29)*.

Lastly, we found that an increase in the induction of the Notch signaling pathway is necessary for VPA to exert its effect. Previously, it was reported that VPA could directly stimulate Notch1 in carcinoid tumors *(7)*. Although we find an effect on the Notch1 receptor, we cannot exclude the involvement of other Notch receptors. Furthermore, additional studies are necessary to investigate whether VPA influences the Notch pathway in a direct or indirect manner.

Taken together, we have been able to optimize a protocol to generate cdMiPs. Furthermore, we found that VPA improves the myogenic capacity of these cdMiPs. This beneficial effect is mediated through an increase in the induction of the Notch signaling pathway. We envision that this novel approach to generate mesodermal progenitors and increase their myogenic potential improves the potential of the cdMiPs to be translated into a form of cell therapy.

## Materials and Methods

### Cell culture and differentiation

#### • iPSCs and cdMiPs

Episomal human iPSCs (ThermoFisher Scientific; Pailsey, UK) and sendai-virus iPSCs were cultured on matrigel-coated plates (Geltrex™ LDEV-Free, ThermoFisher Scientific) in Essential 8™ medium (ThermoFisher Scientific) and maintained under normoxic conditions in a humidified incubator.

Differentiation towards cdMiPs was based on the first four days of a myogenic differentiation protocol, previously described by Shelton et al. *(30)*. Briefly, one hour prior to differentiation, iPSCs were incubated with RevitaCell™ Supplement (100X, ThermoFisher Scientific). Afterwards, PSCs were seeded single-cell on Corning^®^ Matrigel^®^ Matrix-coated plates (VWR, Leuven, Belgium) at a density of 50,000 cells/cm^2^. The following day, differentiation was induced by Essential 6^TM^ Medium (ThermoFisher Scientific) supplemented with CHIR99021 (8μM, Axon Medchem, Groningen, Nederland) for two days. Afterwards, cells were cultured in Essential 6^TM^ medium for another two days. Small molecules such as VPA (1mM, Sigma, Saint-Louis, MO, USA) or DAPT (20μM, Tocris, Bristol, UK) were added from day 2-4. Prior to *in vivo* injections, cells were exposed to 20% FBS for the last 24 hours of differentiation.

#### • Co-culture with C2C12 myoblasts and rat cardiomyocytes

C2C12 myoblasts (Sigma) were maintained under hypoxic conditions in a humidified incubator on collagen-coated plates (Sigma). C2C12 growth medium consisted of Dulbecco modified Eagle’s minimal essential medium (DMEM, 1X, 4.5 g/L D-glucose, L-glutamine) supplemented with 10% FBS, 1% penicillin-streptomycin (pen-strep) and 1% sodium pyruvate, all from ThermoFisher Scientific.

Co-culture with cdMiPs was performed in a 5:1 ratio (cdMiPs:C2C12) on collagen-coated plates (Sigma). The next day, differentiation was induced through culture in DMEM (1X, 4.5 g/L D-glucose, L-glutamine) with 2% horse serum (HS) and 1% pen-strep for five days.

Neonatal rat cardiomyocytes were harvested from new-born rat pups within 24 hours after birth (P179/2013). The minced hearts were digested in ADS buffer (tissue culture water supplemented with 0.0085% NaCl, 0.0005% KCl, 0.00015% monohydrated NaH_2_PO_4_, 0.00125% monohydrated MgSO_4_, 0.00125% glucose, and 0.00595% HEPES) supplemented with 0.4% collagenase II and 0.6% pancreatin (all from Sigma). Herein, the hearts were incubates three times for 20 minutes in a 37°C-warmed water bath, with continuous shaking. Afterwards, the rat cardiomyocytes were purified using a Percoll gradient (0.458 g/ml to 0.720 g/ml; GE Healthcare, Chicago, USA). Cocultures with cdMiPs were performed in a 10:1 ratio (cdMiPs:cardiomyocytes) in high-glucose DMEM supplemented with 18% medium 199, 5% HS, 5% FBS, 1% pen-strep, and 1% L-glutamine (all from ThermoFisher Scientific) on gelatin-coated plates (Bio-connect, Huissen, Nederland).

#### • Differentiation towards mesodermal lineages

For both smooth muscle and osteogenic differentiation, 30,000 cells/cm^2^ were seeded single-cell on collagen-coated plates. To induce smooth muscle differentiation, cells were put in DMEM (1X, 4.5 g/L D-glucose) supplemented with 1% pen-step, 1% L-Glutamine, 1% sodium pyruvate, 2% HS (all from ThermoFisher Scientific) and transforming growth factor beta 1 (TGFβ1) (50ng/ml; Peprotech, New Jersey, USA). For osteogenic differentiation, cells were exposed to αMEM supplemented with 10% FBS, 1% L-Glutamine, 1% pen-strep (all from ThermoFisher Scientific), 50μM Ascorbic-2-phosphate, 10mM β-glycerolphosphate and 10mM dexamethasone (all from Sigma). Full myogenic differentiation of the cdMiPs was performed according to the protocol published by Shelton et al. *(30)*.

### Generation and validation reporter line

The plasmid contained a puromycin resistance cassette (puro), enhanced GFP (eGFP), firefly luciferase (Fluc), and the human sodium iodide symporter (hNIS). Expression was driven by the cytomegalovirus (MLV) early enhancer element together with the chicken beta-actin promoter (CAGGS-promoter). The plasmid was integrated via ZFNs downstream of exon 1 of the PPP1R12C gene on chromosome 19. Here, iPSCs were resuspended in nucleofection solution 2 (Amaxa; Lonza) with 10 μg of donor plasmid and 3 μg of ZFN messenger RNA per 2 × 10^6^ cells. Integration was performed through electroporation (Program F16, Amaxa). After a week of puro selection, individual clones were expanded.

To check for reporter gene incorporation, genomic DNA was extracted using the PureLink™ Genomic DNA Mini Kit (Thermo Fisher), following the manufacturer’s protocol. A PCR was performed to check for the 3’ junction (TTCACTGCATTCTAGTTGTGG and AAGGCAGCCTGGTAGACA) and 5’ Random integration (GTACTTTGGGGTTGTCCAG and TTGTAAAACGACGGCCAG). For Southern Blot analysis, 5μg of genomic DNA was digested with NcoI (New England Biolabs, Massachusetts, US) and loaded onto a 0.7% agarose gel. Fragments were transferred to a nylon membrane (Zeta-Probe; Biorad) that was ultraviolet-crosslinked, and a pre-hybridization was performed. The probe targeting the homology arm was labelled using the Ladderman Labelling kit (TaKaRa, Shiga, Japan) by PCR using the donor vector. Hybridization of the membrane with the probe was performed using the ExpressHyb™ Hybridization solution (TaKaRa).

#### • *In vitro* tracer uptake experiment

iPSCs were plated on a 24-well plate under normal growth conditions. Cells were rinsed and incubated for one hour with 250 μl of ^99m^TcO_4_^-^ (0.74 MBq/ml in DMEM; ThermoFisher Scientific). A blocking experiment was performed with the addition of NaClO_4_ (0.74 MBq/ml ^99m^TcO_4_^-^ in 10 μM NaCl_4_ + DMEM). Afterwards, cells were washed thrice with ice-cold phosphate-buffered saline (PBS; ThermoFisher Scientific), thereby collecting the supernatants. Next, cells were lysed using Reagent A100 lysis buffer and Reagent B Stabilising Buffer (ChemoMetec A/S, Allerød, Denmark) and collected separately from the supernatant. Radioactivity of both the supernatant and the cells was measured by the 2480 Wizard2 Automatic Gamma Counter (PerkinElmer, Waltham, MA, USA). Uptake values were adjusted for tracer decay and corrected for cell amount.

#### • *In vitro* Bioluminescent imaging

iPSCs were exposed to 0.3 mg/l D-luciferin (Promega, Benelux, Leiden, The Nederlands). Images were obtained immediately after D-luciferin application with the IVIS^®^ Spectrum (Caliper Life Science, Hopkington, MA, USA). Photon flux (p/s) was measured and images were further analyzed with Living Image version 4.2 (Caliper Life Science).

### Integration doxycycline-inducible DLL1

A doxycycline (dox)-inducible *DLL1* was integrated by means of RMCE. The master cell line (H9) used for RMCE was generated previously using ZFN-mediated integration of a flippase (FLP) recombinase target-flanked donor cassette into the AAVS1 locus *(14)*. RMCE was performed by nucleofection of the master cell line with a donor vector and the FLPe-expressing vector. Nucleofection was done on 3 x 10^6^ cells obtained after accutase treatment using the hESC Nucleofector Solution Kit 2 (Amaxa) and program F16 using an Amaxa nucleoporator. Donor plasmids were generated through Gibson assembly (NEB) of PCR-amplified open reading frames of the desired genes. Plasmids were then evaluated by digestion and Sanger sequencing. Overexpression of Dll1 was achieved by the addition of 2μg/ml dox.

### Gene expression analysis

Total RNA was extracted using the PureLink^TM^ RNA Mini Kit (Invitrogen) according to the manufacturer’s protocol. Afterwards, the RNA was treated with the DNA-free^TM^ Kit (Invitrogen) and 500ng of RNA was reverse transcribed into cDNA with the Superscript^®^ III Reverse Transcriptase First-Strand Synthesis SuperMix (Invitrogen). Quantitative real-time PCR (qPCR) was performed with the Platinum™ SYBR^TM^ Green qPCR SuperMix-UDG (Invitrogen). The qPCR cycle was performed for two minutes at 95°C, 40 cycles of 15 seconds at 95°C and 45 seconds at 60°C. All primers are listed in Table 1. The obtained Ct values of the tested genes were normalized to the geometric mean of the Ct values of housekeeping genes *RLP13A, GAPDH* and *ACTB*.

**Table 1.**
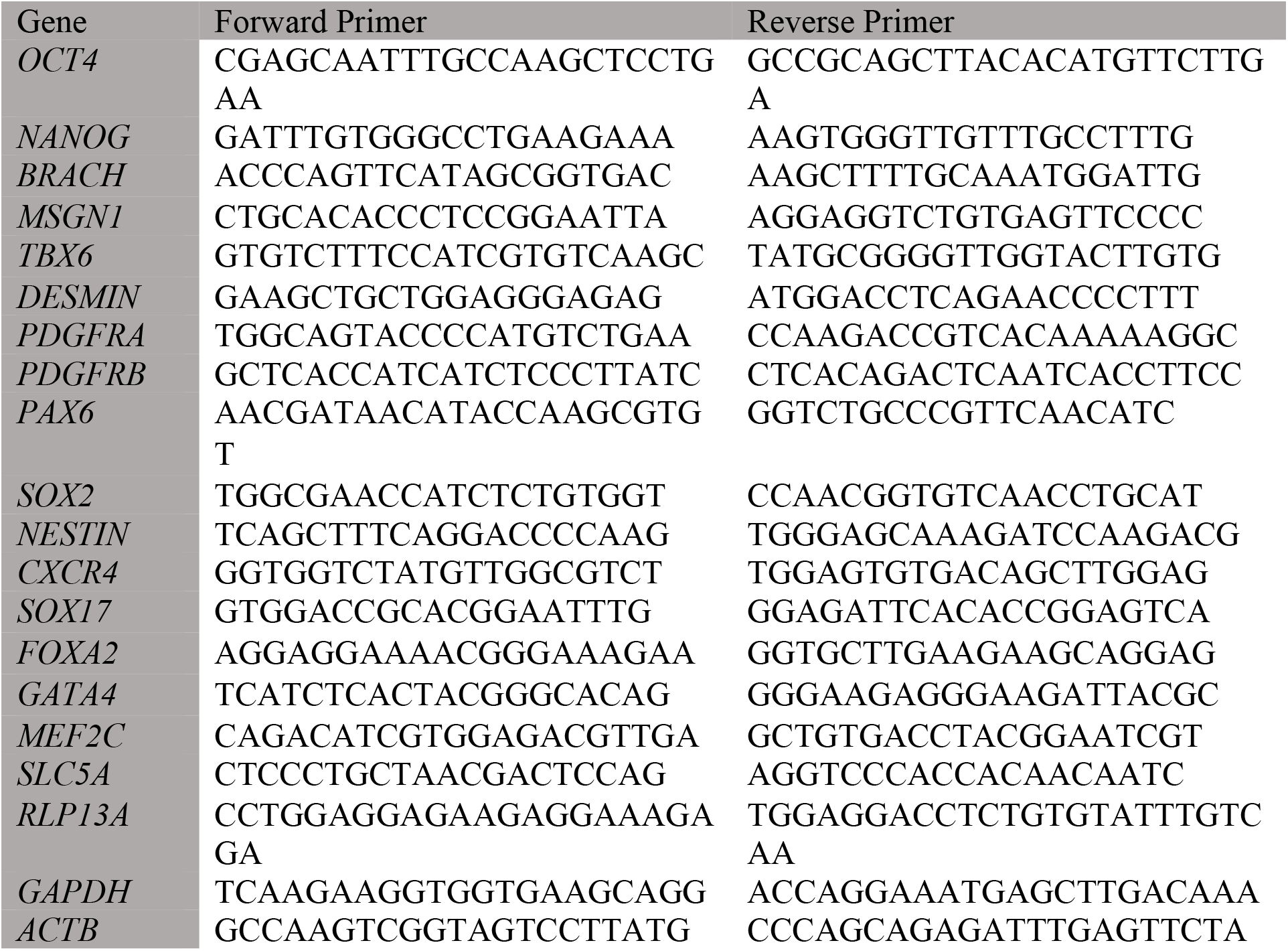
RT-PCR primers from 5’-3’

### Immunostaining

Cells were fixed with 4% paraformaldehyde (Sigma) for 15 minutes at room temperature. Afterwards, cells were permeabilized with 0.02% Triton (Triton X-100; Sigma) in 1% bovine serum albumin (BSA; Sigma) in PBS (Thermo Fisher Scientific) for 30 minutes, followed by a 60-minute blocking with donkey serum (1:10 diluted in 1% BSA; VWR, Pennsylvania, USA). Incubation with the primary antibody was performed overnight at 4°C. The following day, cells were incubated for one hour with AlexaFluor^®^-conjugated donkey secondary antibodies (1:500, Thermo Fisher Scientific) and finally counterstained with Hoechst 33342 (20 μm, Thermo Fisher Scientific). Primary antibodies used were: goat anti-OCT4 (1:200; ab27985; Abcam, Cambridge, UK), rabbit anti-NANOG (1:200; ab80892; Abcam), mouse anti-TRA1-60 (1:50; sc-21705; Santa Cruz, Dallas, USA), mouse anti-MyHC (1:20, Hybridoma Bank), rabbit anti-LAMIN A/C (1:300, ab108595, Abcam), rabbit anti-cardiac MyHC antibody (1:100, ab50967, Abcam), rabbit anti-LAMININ (1:300, L9393, Sigma) and mouse anti-sarcomeric alpha-actinin (SMA; 1:100, ab9465, Abcam). Staining with anti-Actin, α-Smooth Muscle – Cy3™ antibody (1:200, C6198, Sigma) was performed for two hours at room temperature. The presence of calcium deposition after osteogenic differentiation was visualized by incubating the cells with alizarin red S (40 mM, A5533, Sigma, pH4.2) for 10 minutes.

Pictures were taken with an Eclipse Ti microscope (Nikon) by means of Image-Pro Plus 6.0 software (Nikon). Image analysis was performed with ImageJ software (NIH).

### Flow cytometry

Cells were diluted to 1 x 10^6^ cells/ml with FACS buffer (2 % FBS, 10 mM HEPES and 10 mM NaN3 at a pH of 7.2, all from ThermoFisher Scientific) and incubated with the primary antibodies for 30 minutes at 4°C. The following antibodies were used: CD44-FITC (0.25 μg/10^6^ cells, 110441-81, Thermo Fisher), CD140a-PE (10 μl/10^6^ cells, 558002, BDBioscience, Eysins, Switzerland) and CD140b-APC (5 μl/10^6^ cells, A15719, Thermo Fisher). Right before analysis, the cells were stained with SYTOX^TM^ green dead cell stain (1μl/ ml) to assess viability. UltraComp eBeads^TM^ compensation beads (Thermo Fisher) were used for compensation. The analysis was performed on a BD FACSCanto™ II HTS (BD Biosciences).

### Western Blot

Cells were lysed using a urea-thiourea buffer (7M urea, 2M Thiourea, 4% CHAPS, 0.1% β-mercaptoethanol and 30mM Tris HCl pH8.5, all from Sigma) supplemented with cOmplete™ Protease Inhibitor Cocktail (1:100, Thermo Fisher) and phosphostop (1:100, Roche, Basel, Switzerland). 30μg of protein was dissolved in sample-loading buffer (5% SDS, 0.2% bromophenol blue, 50% glycerol and 250mM Tris HCl pH8.5, all from Sigma), loaded onto 10% SDS-polyacrylamide gels and subsequently transferred onto nitrocellulose membranes (Protran, Sigma). Membranes were blocked with Tris-buffered saline (TBS) containing 0.05% Tween and 5% skim milk powder (Sigma) and incubated with the primary antibody overnight at 4°C. The following day, membranes were incubated with the secondary antibody for one hour at room temperature. All secondary horseradish peroxidase-conjugated antibodies (BioRad) were diluted 1:5000 in TBS-Tween and 2.5% skim milk powder. The polypeptide bands were detected with Gel Doc^TM^ chemiluminescence detection system (BioRad) after incubation with SuperSignal™ West Pico Chemiluminescent substrate (Thermo Scientific, Catalog #34087) or SuperSignal™ West Femto Maximum Sensitivity substrate (Thermo Fisher Scientific, Catalog #34095). Relative densitometry was obtained by normalizing the protein band versus background and a housekeeping protein. The primary antibodies used were rabbit anti-Dll1 (0.5μg/ml, PA5-42902, Thermo Fisher) and rabbit anti-Cleaved Notch1 (1:500, 4147, Cell Signaling).

### In vivo differentiation potential of cdMiPs

*Sgcb^-/-^ Rug2γc^-/-^* mice were injected intra-muscularly with either saline or 1.5 x10^6^ cdMiPs (with or without FBS) in the quadriceps, gastrocnemius and tibialis anterior. For the cardiac injection, 1.5 x 10^6^ cells were suspended in 30μl Reduced Growth Factor Basement Membrane Matrix 1:1 diluted in DMEM-F12 and injected in the left ventricle wall. Afterwards, the mice were monitored through bioluminescence imaging (BLI). For *in vivo* BLI scans, mice were placed in the flow chamber of IVIS^®^ Spectrum. Subsequently, 126 mg/kg of D-luciferin was injected subcutaneously. Next, consecutive frames were acquired until the maximum signal intensity was reached. Animal protocols were approved by the Ethical Committee for Laboratory Experimentation, project number P004/2017 and P182/2018. The grafted muscles were harvested four weeks after transplantation, embedded in OCT and snap frozen in liquid nitrogen. Serial transverse 8μm cryostat sections were obtained from cell-injected muscles using the cryostat (Leica, Wetzlar, Germany).

### Single-cell RNA sequencing

CdMiPs were sorted single-cell by FACS in 96-well plates in RLT buffer. mRNA was isolated using the genome and transcriptome sequencing protocol *(31)*. Prior to isolation, external RNA controls consortium (ERCC) spike-in RNAs can be added to the RLT buffer. Further processing was done using the Smart-seq2 protocol *(15)*. Briefly, cells were incubated for 3 minutes at 72 °C. Afterwards, the RNA was reversely transcribed and amplified via PCR for 25 cycles. Amplification was done with KAPA HIFI Hot Start ReadyMix (KAPA Biosystems, Wilmington, USA) and purified by magnetic beads (CleanNA). Quantity and quality of cDNA were assessed with a Qubit fluorometer (Thermo Scientific) and Agilent 2100 BioAnalyzer, respectively. The libraries were prepared with the Nextera XT library prep and index kit (Illumina). 100pg of cDNA was tagmented by transposase Tn5 and amplified with dual-index primers (i7 and i5, 14 cycles). In total 279 single-cell libraries were pooled together and sequenced single-end 50bp on a single lane of a HiSeq4000 (Illumina).

In the Fastq files, tags were trimmed with cutadapt 1.5 *(32)*. The retained tags were aligned to the Ensembl GRC38p7 human reference genome using STAR 2.4.0 *(33)*. Quality control on aligned and counted reads was done with Scater 1.8.4 *(34)*, cells with less than 150,000 reads; less than 2,500 or over 8,500 detected genes; less than 6% mitochondrial DNA and less than 10% of spikein ERCCs were removed. Read normalization was done with Scran 1.8.4. The first ten principal components were used to reduce the dimensionality of the data. Afterwards, t-SNE were computed with a perplexity of 5 in the package Seurat v2.1.0. Clustering was performed with SC3 1.8.0 using k = 2-3. Differential expression between clusters was calculated using a likelihood ratio test corrected for multiple testing with a Benjamini-Hochberg False Discovery Rate correction with MAST. Marker genes defined as genes with areas under the receiver operating curve (AUROC) > 0.85 and with P < 0.01 using SC3 1.8.0 *(35)*.

### Statistical analysis

Statistical analysis and generation of graphs were performed on GraphPad Prism 7.0 (GraphPad Software, San Diego, CA, USA). Two-tailed t test or one-way ANOVA were used to compare interrelated samples, with a Tukey’s post-hoc test to correct for multiple comparison. Two-way ANOVA was used for comparing multiple factors. Data is represented as mean±standard error of the mean (SEM). The number of independent experiments or biological replicates are represented in the figure legends.

## Supplementary Materials

Fig. S1. Integration of imaging reporter genes into a safe-harbor locus.

Fig. S2. Serum pre-exposure does not influence lineage markers.

Fig. S3. Quality control single-cell RNA-sequencing of mesodermal progenitors.

## Acknowledgments

NB would like to thank Jordi Camps and Jens van Herck for the help in performing the SMART-Seq2 and the Master students Zeger Derynck and Laura Danti for their help in some of the experiments. Furthermore, a big thanks to Prof. Olivier Pourquié who kindly provided the MSNG1-Venus reported cell line.

## Author contributions

N.B.: Conceptualization, Methodology, Formal analysis, Investigation, Writing – Original Draft and Visualization. N.G. and L.Y.: Writing – Review & Editing, Validation. P.C.: Methodology, Writing – Review & Editing. S.H.: Resources and Writing – Review & Editing. A.R.: Resources and Writing – Review & Editing, C.D.: Supervision, Resources and Writing – Review & Editing. M.S.: Supervision, Conceptualization, Funding acquisition and Writing – Review & Editing.

## Supplementary Materials

**Fig. S1.**
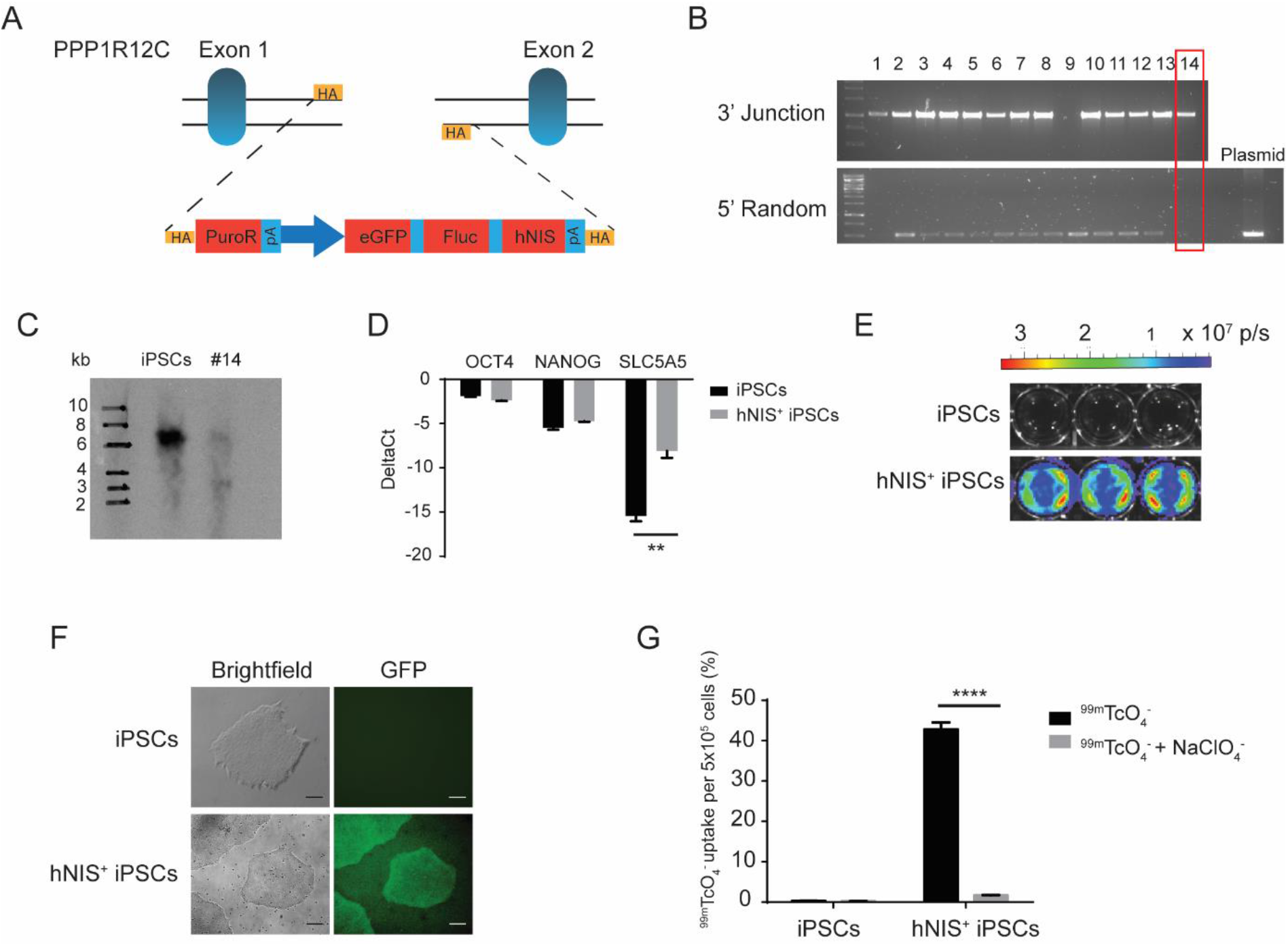
Integration of imaging reporter genes into a safe-harbor locus. (**A**) Schematic representation of the reporter gene construct integrated between exon 1 and 2 of the PPP1R12C gene of induced pluripotent stem cells (iPSCs). (**B**) Junction and random integration assay of the picked clones after zinc finger targeting. (**C**) Southern blot analysis of the selected clone 14 compared to the negative control. (**D**) qPCR of pluripotency markers OCT4 and NANOG and for the human sodium iodide symporter (SLC5A5/hNIS) in the hNIS^+^ line compared to the unedited iPSCs (n=3). (**E**) *In vitro* bioluminescence imaging (BLI) of hNIS^+^ iPSCs compared to hNIS^-^ iPSCs. (**F**) Immunofluorescence showing enhanced green fluorescent protein (eGFP) fluorescence in hNIS^+^ iPSCs compared to hNIS^-^ iPSCs. (**G**) ^99m^TcO_4_^-^ uptake experiment in hNIS^+^ compared to hNIS^-^ iPSCs. NaClO_4_ was used to specifically block the hNIS from taking up ^99m^TcO_4_^-^. Scale bars: 100μm. **P<0.01; ****P<0.0001.

**Fig. S2.**
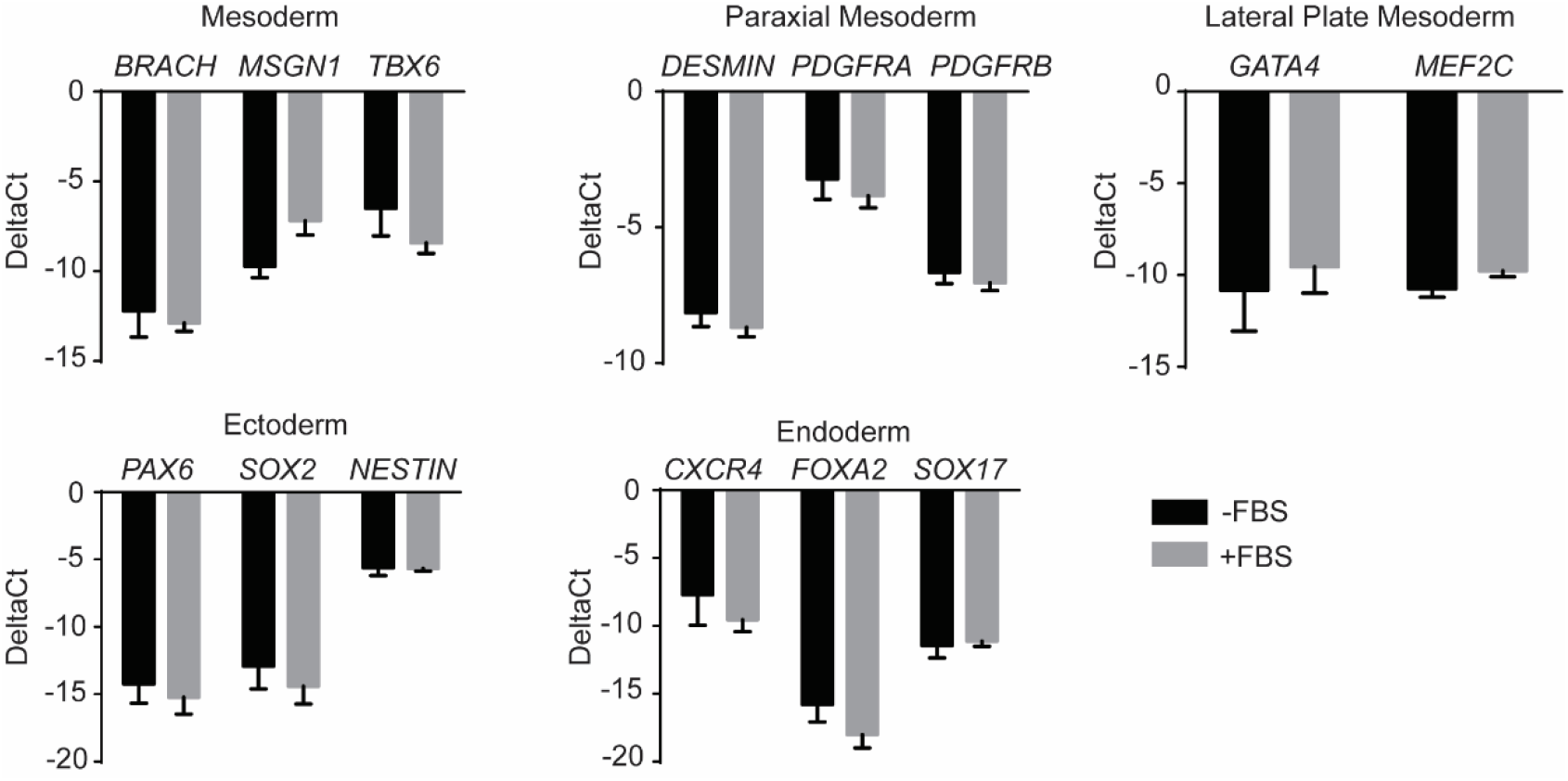
Serum pre-exposure does not influence lineage markers. Gene expression analysis of early and late mesodermal markers as well as markers for ectoderm and endoderm in mesodermal progenitors with or without fetal bovine serum (FBS) pre-exposure (n=3).

**Fig S3.**
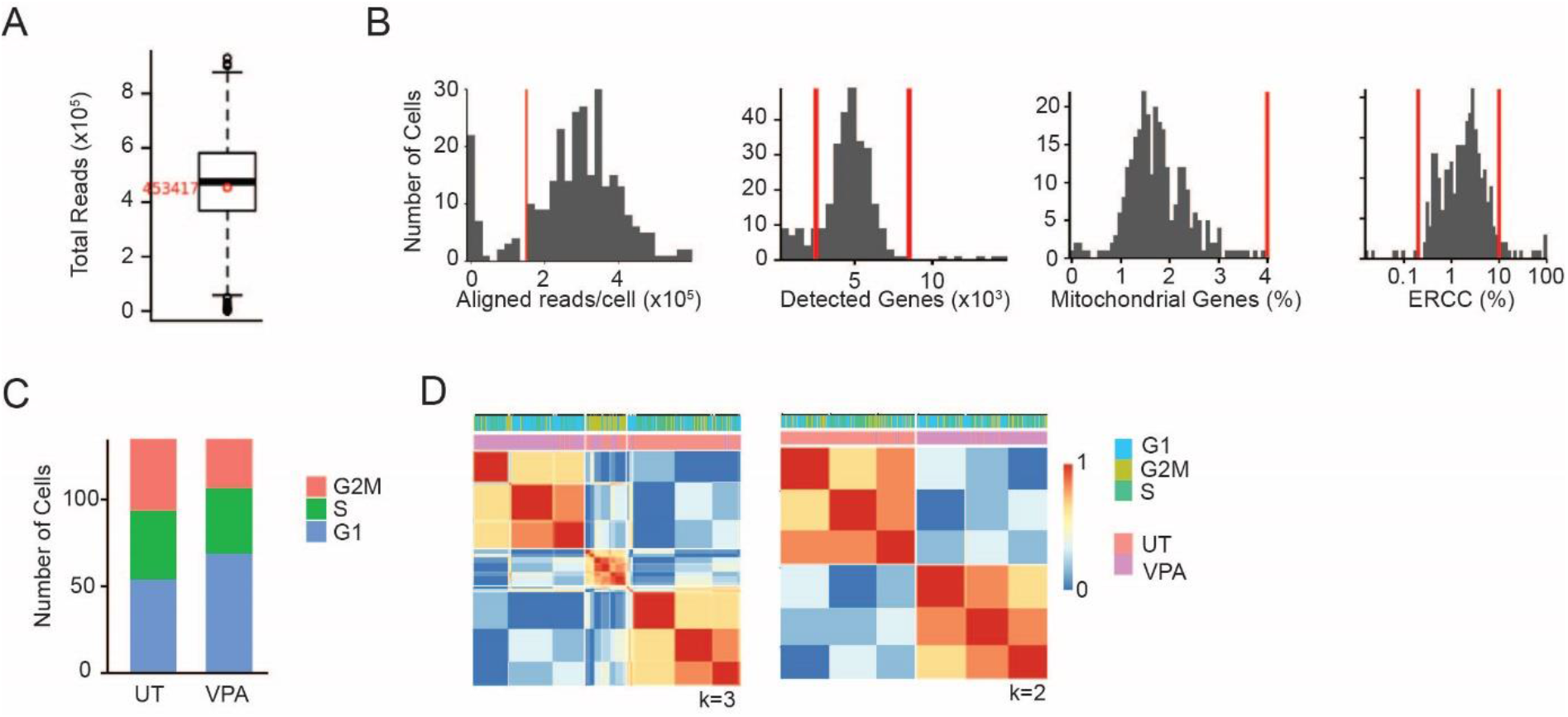
Quality control single-cell RNA sequencing of mesodermal progenitors. (**A**) Boxplot of the aligned reads/cell. (**B**) Quality control of scRNAseq data with cut-offs for aligned reads/cell, detected genes per cell, percentage of mitochondrial genes and percentage of external RNA controls consortiums (ERCCs). (C) Cell-cycle phase for mesodermal progenitors either untreated (UT) or treated with valproic acid (VPA). (**D**) Consensus matrix for k=2-3. Clustering was compared with both cell cycle and treatment condition.

